# Synchronous Miocene radiations and geographic-dependent diversification of pantropical *Xylopia* (Annonaceae)

**DOI:** 10.1101/2025.05.21.655441

**Authors:** Francis J. Nge, David M. Johnson, Nancy A. Murray, Laura Holzmeyer, Keegan Floyd, Gregory Stull, Vincent Soule, Pierre Sepulchre, Delphine Tardif, Carlos Rodrigues-Vaz, Thomas L. P. Couvreur

## Abstract

**Aim:** The evolutionary drivers of hyperdiversity in tropical rain forests are complex and multifaceted. We used the pantropical *Xylopia* (Annonaceae) genus to address the diversification of rain forest lineages through time, across different regions, and into novel non-rain forest habitats with a comparative phylogenetic approach.

**Location:** Global (pantropical)

**Methods:** We generated a time-calibrated phylogeny of *Xylopia* using hybrid capture sequence data, including 88% (168/191 spp.) of species within the genus. Diversification analyses were conducted to test for the presence of rate heterogeneity (BAMM, ClaDS, CoMET) and environmental-dependent (RPANDA), geographic-dependent, and habitat dependent (GeoHiSSE) diversification in *Xylopia*.

**Results:** Significant diversification rate heterogeneity was detected along the backbone of the core *Xylopia* clade, leading to near synchronous radiations across tropical regions globally in the Miocene, with higher diversification rates in Africa, Central America, and Madagascar, and lower rates in Australia + New Guinea. Transitions from rain forest to subhumid habitats led to lower diversification rates, whereas transitions to ultramafic habitats lead to higher rates. Regional diversification models indicate sea-level changes as an important driver for Asian, Australian, Pacific, and Neotropical clades and regional temperature changes as the main diversification driver for an African clade of *Xylopia*.

**Main conclusions:** Our study shows that despite synchronous radiations across regions, different regional environmental drivers have affected the diversification of *Xylopia* across tropical regions globally. A noteworthy example includes radiations of all five Malagasy clades *c.* 7 Ma coinciding with the establishment of heavy seasonal rainfall linked with the Indian monsoon. The diversification dynamics of rain forests are complex and heterogenous, with different clade-dependent and region-dependent environmental drivers.

## Introduction

Tropical rain forests are the most diverse terrestrial biome in the world. Covering just under 10% of land, they contain nearly half of plant diversity (Eiserhardt *et al*. 2017). How this spectacular plant diversity arose is a topic of great interest in evolutionary biology. Some studies show that rain forest plant lineages have diversified at more or less a constant rate, with a steady accumulation of diversity through time based on molecular phylogenies (Couvreur *et al*. 2011a; Feldberg *et al*. 2014; Schneider and Zizka 2017; Bruun-Lund *et al*. 2018; Gamisch and Comes 2019). Other rain forest plant groups such as Malpighiales and Menispermaceae showed an early burst in diversification (i.e., ancient radiations; Davis *et al*. 2005; Wang *et al*. 2012), while others conversely have much younger radiations (Richardson *et al*. 2001; Erkens *et al*. 2007; Särkinen *et al*. 2007; Koenen *et al*. 2015). Heterogeneity in extinction rates and evidence (or absence) of mass extinction events have also been shown in numerous rain forest plant lineages from molecular phylogenies (Meseguer *et al*. 2018; Brée *et al*. 2020; Dagallier *et al*. 2023a). It is clear that there are different macroevolutionary pathways for hyperdiversity in rain forest lineages. However, whether one of these diversification modes is the predominant route to hyperdiversity remains inconclusive.

Large-scale environmental changes have affected tropical regions differently. For example, greater aridification across Africa from the Miocene onwards has been suggested as the main driver for higher extinction rates of tropical lineages on that continent compared to other regions (the odd man out hypothesis; Couvreur 2015). In the Neotropics, a significant portion of plants included in a meta-analysis by Meseguer *et al*. (2022) showed time-dependent diversification, followed by temperature-dependent diversification. Time dependency in determining diversification rates and species richness points to a potentially general universal explanation for biodiversity patterns (Henao Diaz *et al*. 2019; Li and Wiens 2019), though this has been debated (Scholl and Wiens 2016). Other environmental factors such as sea level, temperature, precipitation and the development or change of particular climatic regimes (e.g., monsoonal regimes) can also been incorporated into diversification models to test for their relative effects in addition to time (Condamine *et al*. 2013; Condamine *et al*. 2017a; Kong *et al*. 2017; Rolland and Condamine 2019).

Environmental changes such as climatic cooling and aridification in the Miocene have caused transitions from humid rain forests to drier habitats for several plant groups (Davis *et al*. 2002; Holstein and Renner 2011; Veranso-Libalah *et al*. 2018; Chen *et al*. 2019; Zizka *et al*. 2020). In nearly all these cases, multiple independent transitions have occurred throughout the evolutionary history of these groups, coinciding with the onset and/or expansion of drier biomes (Simon *et al*. 2009; Vasconcelos *et al*. 2020; Couvreur *et al*. 2021). In some cases, these transition events are linked to increases in diversification rates and radiation into drier habitats (Bouchenak-Khelladi *et al*. 2010), while others are not (Couvreur *et al*. 2011c; Veranso-Libalah *et al*. 2018; Zizka *et al*. 2020). Thus, biome transitions out of rain forest do not always lead to subsequent diversification as commonly thought (Donoghue and Edwards 2014).

These findings on macroevolutionary dynamics of rain forests may change as we get higher taxonomic sampling and phylogenetic resolution resolution. For example, Annonaceae was shown to have a constant diversification rate based on a genus-level phylogeny (Couvreur *et al*. 2011b). However, with greater sampling at the species level, several recent diversification rate shifts in the family are evident, mostly for species-rich genera (Xue *et al*. 2020). Indeed, concentrated sampling for some of these genera has also revealed recent radiations such as for *Guatteria* (Erkens et al. 2007), Monodoreae tribe (Dagallier *et al*. 2023a), and *Piptostigma* (Brée et al. 2020). A similar pattern was observed for palms with an overall constant diversification rate across genera (Couvreur *et al*. 2011a), but with several recent increases in diversification rates inferred within genera (Baker and Couvreur 2013; Cano *et al*. 2022; Kuhnhäuser *et al*. 2025). Many of these radiations are not pantropical and are limited to one or two regions only.

In this study, we examine the diversification of rain forest lineages through time, across different regions, and into non rain forest habitats by focusing on the species-rich pantropical genus *Xylopia* (Annonaceae). The genus contains *c.* 191 species distributed across the tropics, with roughly equal species richness among the Neotropics (c. 56 species), Africa and Madagascar (c. 78 species), and Asia–Pacific (c. 57 species). The genus is also present in well-known tropical biodiversity hotspots *sensu* Myers et al. (2000) such as Madagascar (30 spp.; Johnson and Murray 2020), New Caledonia (4 spp; Johnson *et al*. 2013), and outlying Pacific islands. Recent comprehensive taxonomic revisions have provided greater confidence in our estimate and knowledge of species richness found across these different regions for *Xylopia* (Johnson *et al*. 2013; Johnson and Murray 2015; Stull *et al*. 2017; Johnson and Murray 2018, 2020; Johnson and Murray 2023). *Xylopia* is the only Annonaceae genus with a pantropical distribution, allowing us to test for differences in diversification rates across regions. Other studies on pantropical genera at the species level such as *Ficus* (Moraceae), *Bulbophyllum* (Orchidaceae), and *Manilkara* (*Sapotaceae*) have shown that species richness disparities across regions were due totime-for-speciation effects (constant diversification) or differences in diversification rates, or even a combination of both (Armstrong *et al*. 2014; Bruun-Lund *et al*. 2018; Gamisch and Comes 2019). Whether similar complexity in diversification trends is also present in *Xylopia* remains to be tested.

Xue *et al*. (2020) detected an elevated rate of diversification for the crown node of *Xylopia* compared with the background diversification rate in Annonaceae, but their study included only 48 species (25%, 48/191 spp.) of the genus. Most of these species were sourced from a previous study on the infrageneric classification of *Xylopia,* which obtained four plastid regions for 44 species and recognised five sections within the genus (Stull *et al*. 2017). In Thomas *et al*. (2015), the stem divergence of *Xylopia* was estimated at *c.* 55 Ma and the crown at 30 Ma in the Oligocene.

While most species in *Xylopia* are confined to tropical rain forests, there are numerous species found in non-rain forest open habitats (Johnson et al. unpublished)(Johnson and Murray 2018, 2020). These include subhumid habitats in Africa, Asia, and South America, and exposed ultramafic substrates in New Caledonia and Cuba. Plants growing in ultramafic environments are specialised to deal with metal toxicity and often accumulate these elements (e.g. nickel) in their tissues (Garnica-Díaz *et al*. 2023). Whether specialisation in these habitats for *Xylopia* represent ‘evolutionary dead ends’ or opportunities for diversification remains to be tested (Day *et al*. 2016).

Here, we present the first densely-sampled, time-calibrated phylogenomic framework for *Xylopia* to address the following questions on tropical rain forest evolution at a global scale: (1) Is there significant diversification rate heterogeneity across lineages of *Xylopia*? (2) Was the assembly and diversification across different tropical regions synchronous or asynchronous? (3) Are there significant differences in diversification rates and environmental correlates across different tropical regions, due to idiosyncratic environmental drivers specific to each region? And finally, (4) did transitions from rain forest to other environments (subhumid and ultramafic) lead to increases in diversification rates?

## Material and Methods

### 2.1 Sampling, DNA sequencing and data processing

We sampled 183 *Xylopia* and six outgroup samples, of which 168 *Xylopia* species were included in our final dataset after excluding samples with poor DNA yields or poor sequence coverage (<50% genes included)(Table S1) This 168 taxa dataset represents 88% of accepted, published *Xylopia* species diversity (168/191 spp.), spanning the taxonomic and geographic breadth of the genus. All sections recognized in *Xylopia* were sampled. Our study included 48/c. 56 species from the Neotropics, 73/c. 78 from Africa/Madagascar, and 47/c. 57 from Asia and the Pacific.

#### DNA sequencing and data processing

Sequencing was performed using an Annonaceae-specific bait kit, which includes probes for 469 nuclear loci (Couvreur et al. 2019). This bait kit has been successful used in several recent studies (Brée *et al*. 2020; Dagallier *et al*. 2023b; Martínez-Velarde *et al*. 2023; Lopes *et al*. 2024; Nge *et al*. 2024). Protocols for lab work including DNA extraction, library preparation, and sequencing follow previous studies (Couvreur *et al*. 2019; Soulé *et al*. 2023; Soulé *et al*. 2024).

Our pipeline for sequence data processing is outlined briefly here. Raw sequences were demultiplexed, trimmed for quality, and sorted into paired reads per sample. The processed reads were then assembled using the HybPiper v.1.2 pipeline (Johnson et al. 2016). We extracted the supercontig outputs from the HybPiper pipeline for subsequent alignment; loci that were flagged as potential paralogs were excluded as paralogous loci from recent duplication events may not be detected though manual examination. This conservative approach to exclude flagged loci from downstream analysis follows the examples of Shee et al. (2020) and Kuhnhäuser et al. (2021). Furthermore, we followed Shee et al. (2020) in filtering out assemblies with poor coverage across the targeted loci and reduced noise by excluding (1) underrepresented (proportion of species/genes for which sequences were obtained), (2) incomplete (proportion of target sequence obtained for each species/gene), and (3) unevenly distributed sequences (how evenly the sequence lengths are distributed across species/genes). For each sample, the overall coverage score was calculated in R by combining the three metrics via a cube root function. Sequences with an overall score below 0.5 were excluded from subsequent analyses. Included sequences were aligned using MAFFT v.7.305 (Katoh and Standley 2013), with each locus aligned separately. We then trimmed the alignments to remove poorly aligned regions using Gblocks (Castresana 2000) with the ‘-b5=a’ parameter to allow for gap positions.

#### Phylogenetic reconstruction and divergence-time analyses

We applied both concatenated (CON) and multi-species coalescent (MSC) approaches for phylogenetic reconstruction. The CON-phylogeny was generated using RAxML v.8.2.9 (Stamatakis 2014) with the GTRGAMMA substitution model and 100 bootstrap (BS) replicates (Abadi et al. 2019). The software ASTRAL v.5.6.3 (Zhang et al. 2018) was used to construct the MSC-phylogeny. Individual gene-trees for each sequenced locus are required as the input files for ASTRAL. These gene-trees were reconstructed, as above, using RAxML with the GTRGAMMA substitution model and 100 bootstrap replicates. To improve signal and decrease phylogenetic noise, nodes with very low bootstrap support (< 10 BS) were collapsed using Newick utilities v.1.6 (Junier and Zdobnov 2010) prior to the ASTRAL analysis as per recommended practice (Zhang et al. 2018). Phylogenetic reconstruction steps above were conducted on a local HPC cluster (Montpellier, France). Conflicts between the CON- and MSC-phylogenies were visualized as a tanglegram using the ‘cophylo’ function from the phytools package (Revell 2012) in R.v.4.0 (R Core Team 2016).

We estimated divergence times for lineages within *Xylopia* using BEAST v.2.6.6 (Bouckaert et al. 2014). To decrease rate heterogeneity across loci and to reduce the computational burden of examining the entire dataset, we selected a subset for the dating analyses. The 30 most clock-like loci (based on root-to-tip variance) were selected using SortaDate (Smith et al. 2018). These loci were concatenated using Phyx v.1.3 (Brown et al. 2017) for the BEAST analyses. We constrained the starting tree topology to that inferred from the full dataset (i.e., all loci excluding paralogs and badly sequenced loci). For successful BEAST initialization, the starting tree (RAxML) was dated using treePL (Smith and O’Meara 2012). The treePL output was exported as a newick format for configuration in beauti v.2 for BEAST.

Beauti v.2.4.7 (Drummond *et al*. 2012) was used to configure the xml file for BEAST. An uncorrelated lognormal relaxed molecular clock model and birth-death tree prior model were chosen. The substitution model was GTRGAMMA (Abadi et al. 2019) without the proportion of invariant sites to avoid overparameterization (Yang 2006). In order to fix the topology throughout the BEAST runs based on our provided starting tree, four operator parameters were removed in beauti: (1) Wide-exchange, (2) Narrow-exchange, (3) Wilson-Balding, and (4) Subtree slide (https://www.beast2.org/fix-starting-tree/).

There are no confirmed fossil records of *Xylopia*. Rásky (1956) reported a fossil fruit or infructescence from the Eocene of Hungary with radiating finger-like structures and highlighted potential affinities with *Xylopia*, naming the fossil *Xylopiaecarpum eocaenicum* Rásky. This fossil has been cited elsewhere (e.g. Trájer 2024), but not subjected to a careful systematic re-evaluation. Unfortunately, the morphological details provided by the fossil—a single compression specimen—are limited and somewhat ambiguous. It is not clear if the structure represents an aggregate fruit or simple fruits from separate flowers. Details of the seeds, which might provide critical evidence given the distinctive nature of Annonaceae seeds (Corner 1949; van Setten and Koek-Noorman 1992), are absent. We therefore cannot accept this as fossil evidence of *Xylopia*, necessitating the use of secondary calibrations for our dating analyses.

We used three secondary calibration points as per recommended best practice (Sauquet et al. 2012). Namely (1) *Xylopia* stem (min–max; 34.8–58.0 Myr), (2) Annonoideae crown excluding *Guatteria* and Bocageeae (min–max; 39.4–60.8 Myr), (3) Annonoideae crown excluding Bocageeae (min–max; 43.5–65.5 Myr). These secondary calibration points were extracted from an unpublished dated phylogenetic tree of Magnoliids based on the data of Helmstetter et al. (2024). Molecular dating was based on five verified and phylogenetically placed fossils from Massoni et al. (2015). A uniform prior was imposed for the three secondary calibrations using the 95% confidence intervals of node age estimates from the backbone chronogram as the minimum and maximum age constraints (Sauquet et al. 2012). Two independent BEAST analyses ran for 400 million MCMC generations each, sampling every 1,000th generation. The first 20% of runs were discarded as burn-in. The output was analysed using Tracer v.1.7.1 (Rambaut et al. 2014) to check for convergence—indicated by an ESS above 200. Trees from the two independent BEAST runs were combined using LogCombiner v.2.4.7 and summarized in TreeAnnotator v.2.4.7. BEAST runs were completed using the CIPRES v3.3 Science Gateway Portal (Miller et al. 2010). The resulting dated phylogeny was used for subsequent comparative analyses.

#### Diversification rate heterogeneity analyses

To test for the presence of significant diversification rate shifts in *Xylopia*, we used the program Bayesian Analyses of Macroevolutionary Mixtures (BAMM; Rabosky 2014) and our dated BEAST phylogenetic tree as input. A global sampling regime of 90% was specified in the script given our sampling of described and accepted *Xylopia* diversity (168/191 spp.). BAMM analyses were conducted for 8 million rjMCMC generations and stopped when convergence was achieved (ESS value > 200; Table S2). Convergence assessment and post-analyses were carried out using BAMMtools v.2.1.10 (Rabosky *et al*. 2014) in R v.3.5.1 (R Core Team 2016), with the first 10% of MCMC chain discarded as burn-in. One expected rate shift is specified for this prior as recommended for small trees (< 500 tips; Rabosky 2014). We also used ClaDS (Maliet and Morlon 2022) from the PANDA package v.0.8 in Julia v1.10 to estimate branch-specific diversification rates. The ClaDS analysis was performed with 10,000 iterations in three independent chains and a global sampling fraction (f=168/191) was applied to compensate for the missing species. Rates-through-time (RTT) plots were also generated from BAMM, ClaDS, and TESS v.2.1.2 (Höhna *et al*. 2016). We also used the compound Poisson process on Mass-Extinction Times model (CoMET; May *et al*. 2016) implemented in the R package TESS to test for potential mass-extinction events over the course *Xylopia*’s evolutionary history. Similar to the other diversification analyses, a sampling fraction of 90% was specified for CoMET. We also ran CoMET with two different survival probability priors across the mass extinction event (low: 20% surviving; and high 50% surviving). The autostop function was applied to stop the CoMET runs automatically after convergence (ESS > 200).

#### Environmental-dependent diversification

We applied paleoenvironmental birth-death models of Condamine et al. (2013) using RPANDA v2.3 (Morlon et al. 2016) in R to test for climate-dependent diversification patterns (H3). *Xylopia* was divided into five clades based on the current distribution with clades occurring on one or more continental regions (Table 1). Distributional information for each clade was sourced from Johnson et al. (unpublished). We first fitted a constant-rate birth-death model to establish a baseline for speciation (null model) (λ) and extinction (μ). We then fitted a set of four time-dependent models (Morlon *et al*. 2011) in which λ(t) and μ(t) could vary through time: BTimeVAR, variable pure birth model; BTimeVARDCST, variable birth and constant death model; BCSTDTimeVAR, constant birth and variable death; BTimeVARDTimeVAR, a variable birth and death model. Next, we tested a suite of environment-dependent models (Condamine et al., 2013) to assess whether diversification rates correlated with environmental variables, including regional temperature (Tardif *et al*. 2025), sea level (Miller *et al*. 2005; Miller *et al*. 2020), and continental fragmentation (Zaffos *et al*. 2017). The environmental variables tested were chosen individually for each clade to ensure ecological and geographical relevance. All models were evaluated using small-sample corrected Akaike Information Criterion (AICc). We computed delta AICc and Akaike weights to select the best-fit model for each clade and to evaluate relative model support. Models were categorized into six groups: Null (constant birth), Time (time-dependent), GTemp (global temperature), regional temperatures (EURATemp – Eurasian temperature, SAMTemp – South American temperature, AFRTemp – African temperature), Frag (continental fragmentation), and Sea (sea level). The weighted AICc values were recalculated for the best fitting model of each category. The tested models were then ranked based on their weighted AICc and the top three models were used as long as they extended over at least 10%.

**Table 1.**
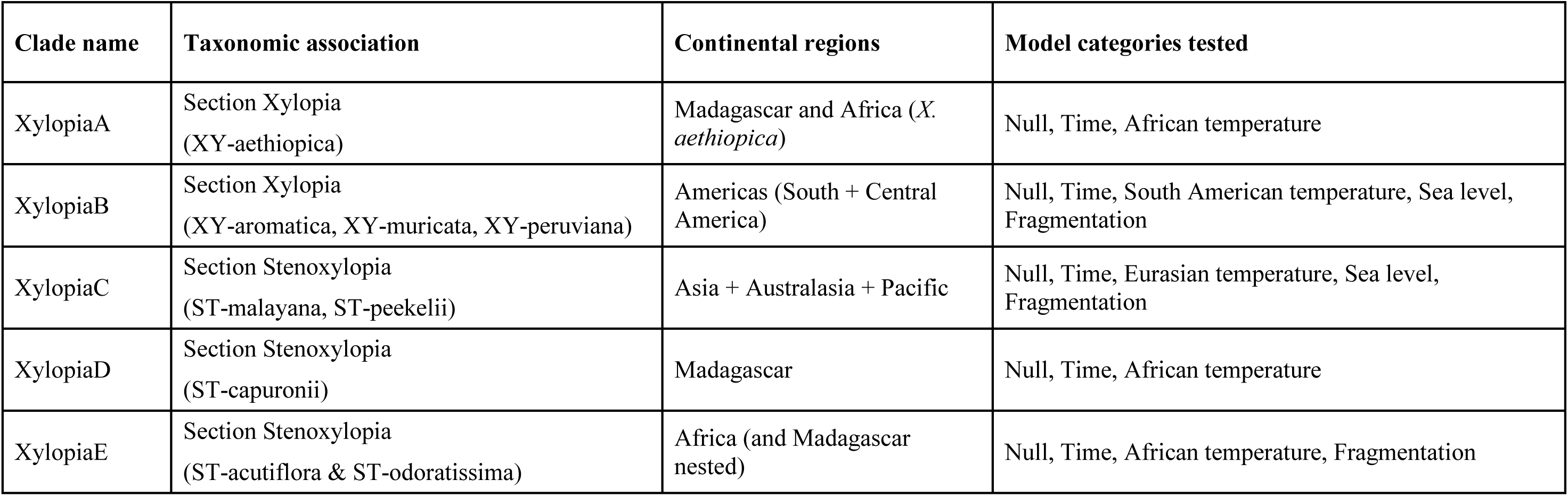
The five *Xylopia* subclades included in RPANDA analyses, to test for environmental-dependent diversification.

#### Geographic and biome diversification

We used GeoHiSSE v.1.9.6 (Caetano *et al*. 2018) in R to compare diversification (speciation-extinction) rates across (i) different geographic regions, (ii) subhumid vs. rain forest (humid) lineages, and (iii) ultramafic vs. non-ultramafic associated lineages. Following Johnson et al. (unpublished) and Chen *et al*. (2019), we defined subhumid environments on the basis of precipitation levels below those typically found in rain forest habitats. We used mean precipitation data to score species as occurring in either humid or subhumid environments. Species with annual mean precipitation lower than 1800 mm and mean precipitation of driest quarter lower than 300 mm were scored as occurring in subhumid environments. Precipitation across different regions were based on climate data from Hijmans *et al*. (2005). In brief, *Xylopia* species occupy subhumid locations in the Sudanian and Zambezian regions of Africa (Linder *et al*. 2012; Johnson and Murray 2018), western Madagascar, and east-central Brazil, as well as in more limited areas in Southeast Asia, Sri Lanka, Cuba, northern Peru, and New Caledonia. GeoHiSSE includes 36 different models that test for geographic-dependent diversification while incorporating hidden states as explanatory variables. As GeoHiSSE only allows for binary traits, geographic coding was conducted in multiple separate analyses for all regions (e.g., Madagascar vs. non-Madagascar). The best fitting model out of the 36 models was determined based on the highest weighted Akaike’s information criterion (AIC) score. A sensitivity analysis was conducted with lineages inhabiting inundated areas coded as rain forest, as they persist in wet-humid localized areas even in subhumid habitats e.g., along riparian vegetation. For ultramafic lineages, both facultative and obligate ultramafic lineages were coded as occurring in ultramafic substrates, vs. other lineages (non-ultramafic). The coding for the geographic regions, subhumid vs. rain forest, inundated, and ultramafic taxa were sourced from Johnson et al. (unpublished).

## Results

### Phylogeny reconstruction and divergence-time estimation

In total, 168 taxa plus outgroups were sequenced successfully, representing ca. 88% (168/191 spp.) of *Xylopia* species diversity. Of the 469 loci sequenced, 197 loci had paralogy issues and were removed (Table S3), resulting in a dataset of 272 and 267 loci for CON and MSC analyses respectively; 5 fewer loci were retained for the ASTRAL analysis due to poor gene recovery of outgroups (rooted gene-trees are required for ASTRAL). The final CON-alignment was 472,273 bp in length, and the subset alignment of the 30 most clock-like loci for the BEAST analysis was 62,780 bp in length.

*Xylopia* was recovered as monophyletic in both our CON- and MSC-phylogenies (Figs. 1, S1). The phylogenies were well resolved, with a strongly supported backbone separating different sections *sensu* Stull *et al*. (2017) and sect. *Rugosperma* as sister to the remainder of the genus. All sections were recovered as monophyletic except that sect. *Verdcourtia* is now nested in sect. *Stenoxylopia* (Fig. 1). Topological conflicts between the CON- and MSC-phylogenies were mostly restricted to shallow regions of the phylogeny, within geographically restricted subclades of the 13 labelled clades (Fig. 1). The backbone topologies from CON- and MSC-datasets were similar except for the placement of XY-muricata which was sister to XY-aromatica for CON but sister to XY-peruviana in MSC, albeit with weak support for the latter. For the CON-tree, only a few nodes were poorly supported (BS < 75); these include the relationship of *X. anomala* with other members of the ST-capuronii clade (BS = 58), the sister relationships of *X. danguyella* and *X. ghesquiereana* (BS = 39), the *X. dibaccata–X. takeuchii* subclade in the ST-peekelii clade (BS = 56), and the position of *X. flexuosa* within the XY-aethiopica clade (BS = 48). Relationships within the ST-capuronii clade are poorly supported in both analyses. Sections *Rugosperma*, *Neoxylopia*, and *Ancistropetala* are all small, geographically restricted depauperate lineages. The bulk of *Xylopia* diversity resides in the *Xylopia* + *Stenoxylopia* sections clade, referred to informally here as the ‘core *Xylopia* clade’. The Neotropical clade (XT-aromatica + XY-muricata + XY-peruviana) is fully supported as monophyletic in both analyses (Fig. 1).

**Fig. 1.**
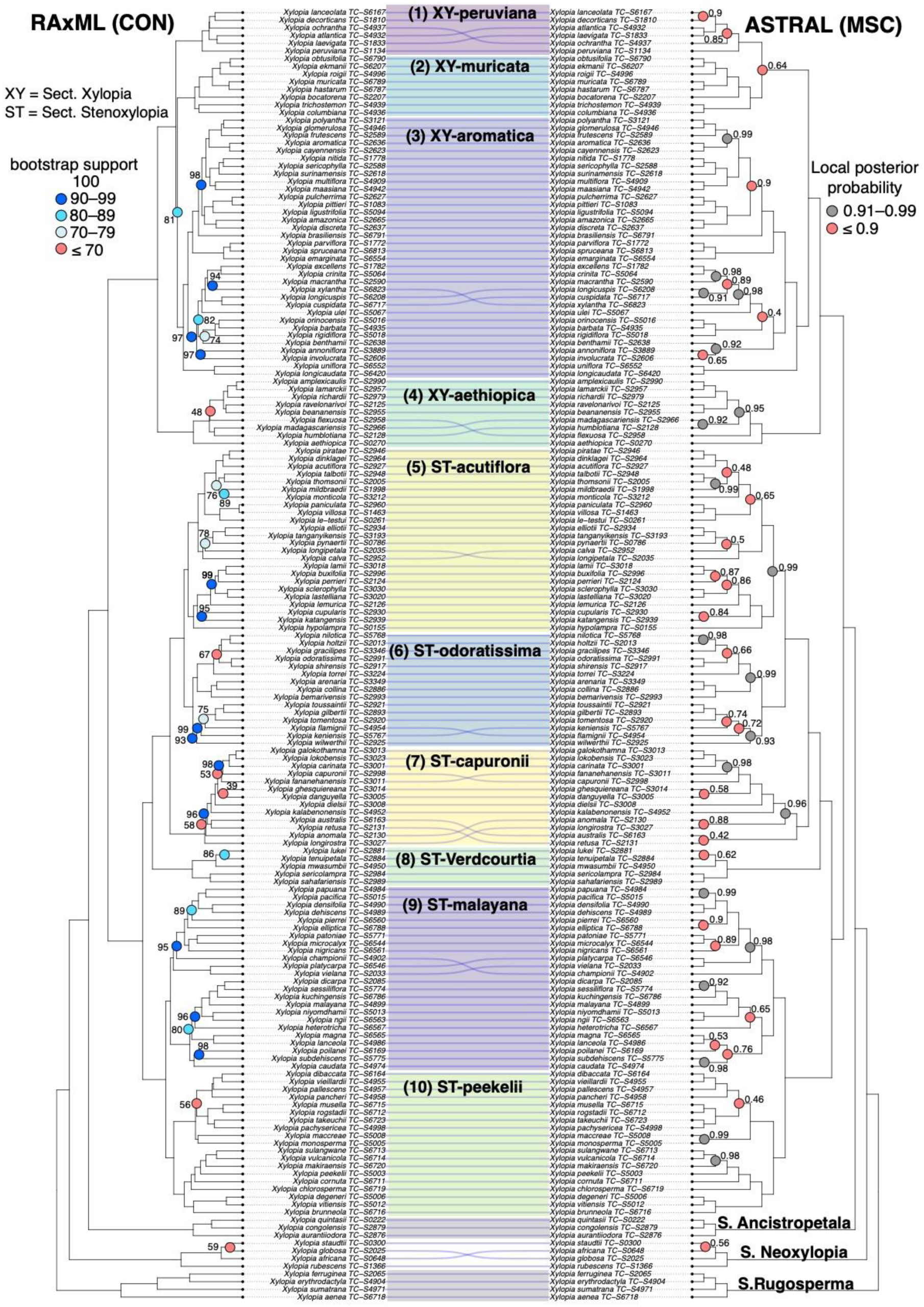
Tanglegram showing phylogenies of *Xylopia* estimated with concatenated (RAxML, left) and coalescent (ASTRAL, right) approaches. For ASTRAL and IQTree, only support values of less than 1 and 100 are shown respectively.

We recovered an Early Oligocene age for crown radiation of the genus (31.77 Ma, 95% confidence interval CI: 22.6–41.02 Ma), based on our BEAST analysis (Table 2, Figs. 2, S2). The earliest divergences (leading to sects. *Rugosperma*, *Neoxylopia*, *Ancistropetala*, and the core clade) occurred in the Oligocene, with the crown age of the core *Xylopia* clade occurring near the Oligocene-Miocene boundary (23.0 Ma, 95% CI 16.2–30.6 Ma). The crown ages of all sections (*Rugosperma*, 20.79 Ma; *Neoxylopia*, 10.54 Ma, *Ancistropetala*, 8.46 Ma; *Stenoxylopia*, 17.24 Ma; *Xylopia*, 14.53 Ma) and sectional subclades were dated to the Miocene, with most of the diversification in the genus (regarding the origins of extant diversity) occurring in the Middle to Late Miocene, after the Mid-Miocene Climatic Optimum (c. 15 to 5 Ma; Steinthorsdottir *et al*. 2021).

**Fig. 2.**
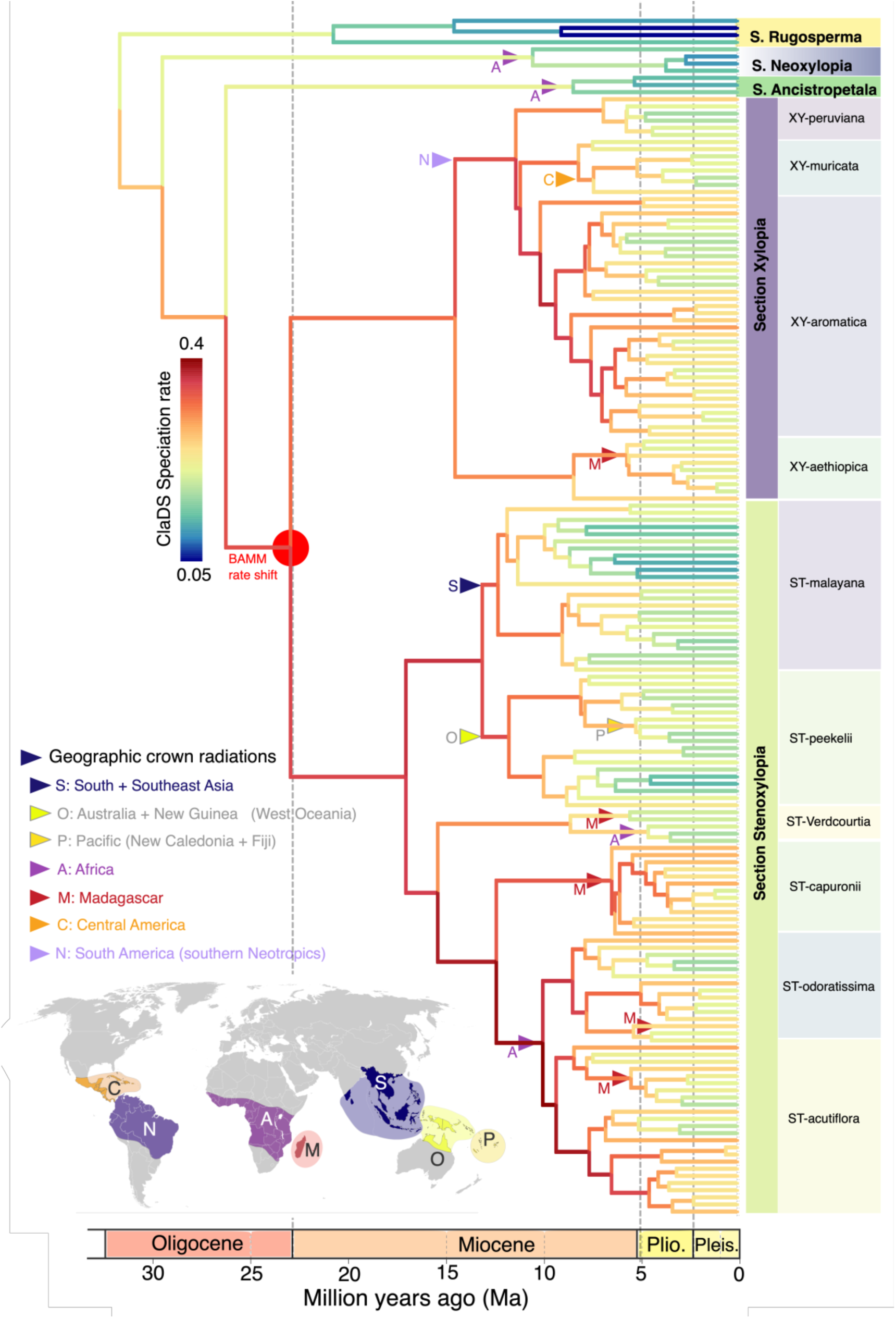
Branch-specific speciation rates for *Xylopia* from ClaDS. The red circle indicates the most probable diversification rate shift from BAMM (at core *Xylopia*: sections *Xylopia* + *Stenoxylopia*). Coloured arrows show the position of crown radiations (≥ 3 spp.) across different geographic regions based on Johnson et al. (unpublished).

**Table 2.**
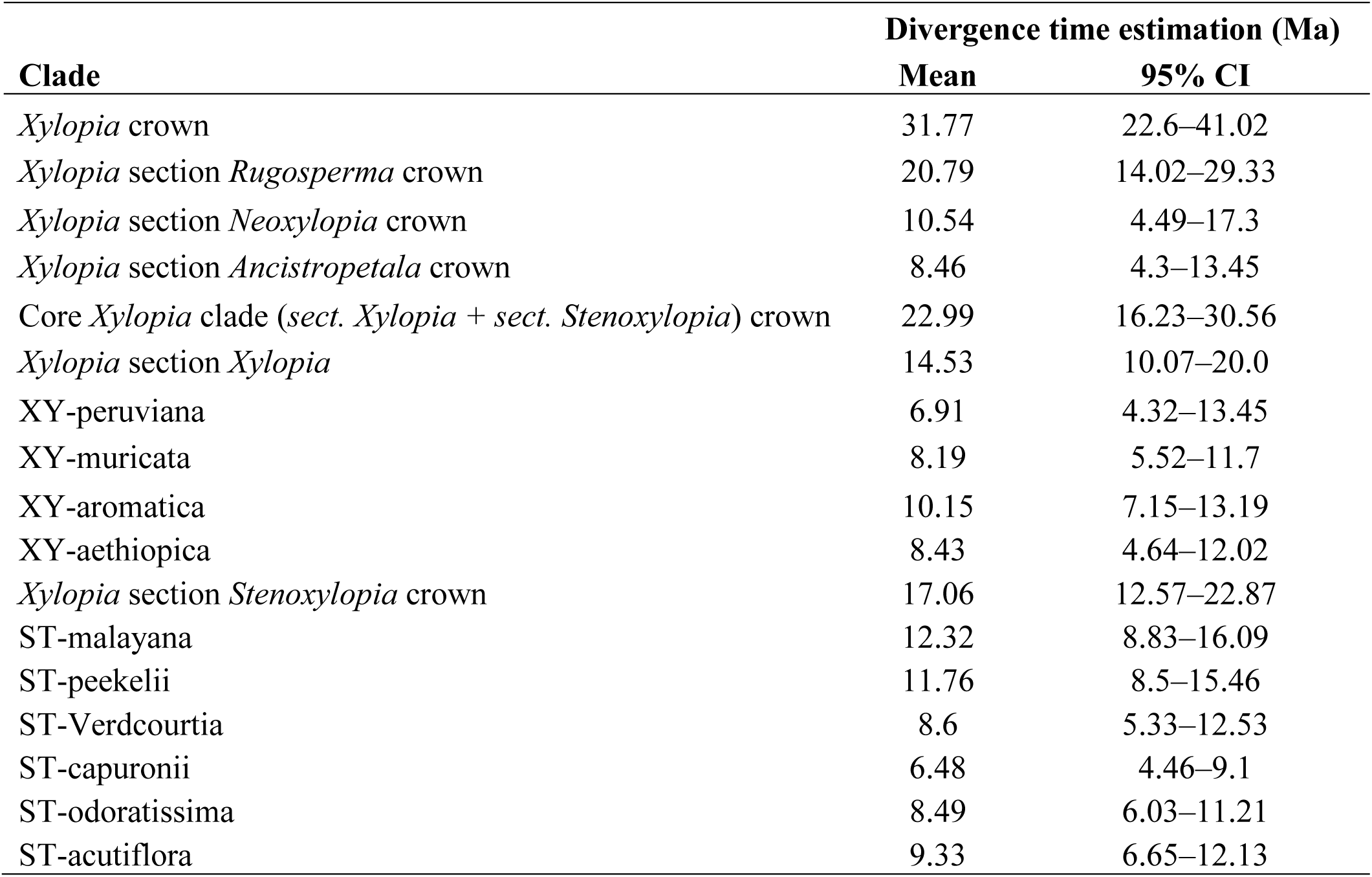
BEAST divergence age estimates of sections and subclades of *Xylopia*.

#### Diversification dynamics

From our BAMM analysis, a single rate shift increase was identified at the crown of the core *Xylopia* clade (Fig. 2, Table S4). The speciation rates-through-time (RTT) analysis from BAMM showed a sharp increase *c.* 23 Ma coinciding with the radiation of the two sections within the core clade around the Oligocene–Miocene boundary. However, following this initial radiation of the core *Xylopia* clade, all subclades across different geographic regions showed a gradual decrease in speciation rates towards the present based on RTT plots (Figs. S3–S7). Similar results were obtained from ClaDS, with highest speciation and diversification rates found along the backbone of *Xylopia* (Fig. 2). Speciation ranges between 0.05 and 0.4 events per million years, with an increase from the root to the highest values distributed in the backbone of the sections *Xylopia* and *Stenoxylopia* subsequently decreasing towards the present. Speciation in the early divergent sections *Rugosperma*, *Neoxylopia*, and *Ancistropetala* does not reach 0.2 events per million years. The modelled extinction rate is close to zero, therefore speciation and diversification rate are nearly equivalent. The Southeast Asian and Pacific *Stenoxylopia* (ST-malayana & ST-peekelii) display lower rates than their sister African/Malagasy clade.

The speciation trend from TESS showed an increase *c.* 15 Ma, followed by a sharp decline from the Pliocene (5.3 Ma) towards the present (Fig. 3C). The extinction rates as inferred from BAMM were quite low, showing a constant decrease through time (Fig 3B), similar to results from TESS with 50% survival probability (Fig. S8). Different results were obtained from TESS with a 20% extinction survival probability, showing a sharp increase in extinction from the Pliocene (5.3 Ma) towards the present (Fig. 3E). No mass extinction events were detected from our CoMET analyses (Figs. 3G,H, S8).

**Fig. 3.**
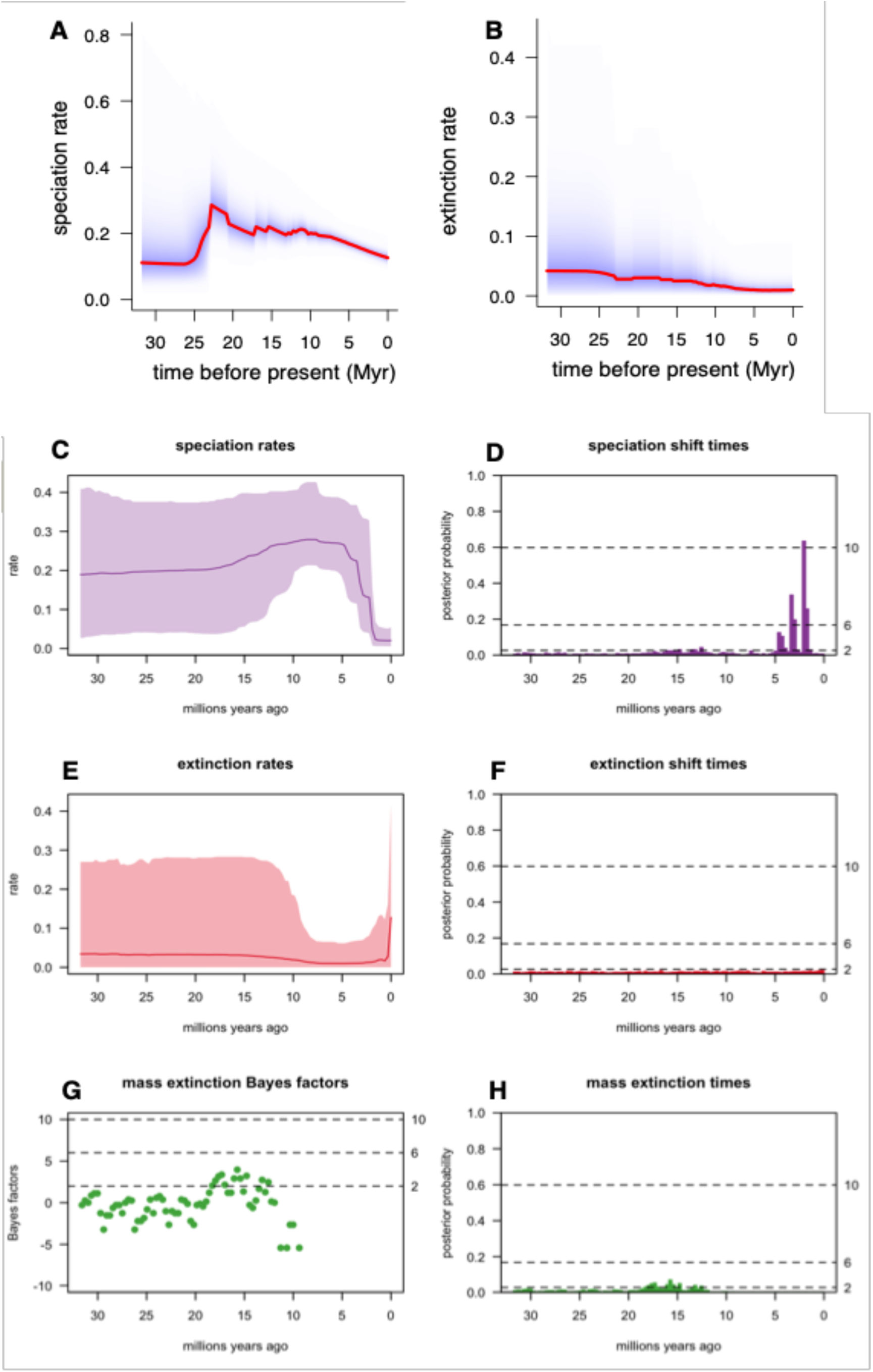
Rates-through-time plots for *Xylopia* from BAMM showing (a) speciation, (b) extinction rates, from TESS showing (c, d) speciation rates and shifts, (e, f) extinction rates and shifts, from CoMET (g, h) probability of mass extinction events through time. These analyses were completed based on the specified prior of 20% survival probability for mass extinction events.

#### Environmental-dependent and regional diversification

Our results returned different favoured environmental-dependent models depending on the regional clades we defined. The best fitting model for *Xylopia* clades B (Neotropics) and C (Asia-Australia-Pacific) was the sea level fluctuation under a pure birth with exponential variation (BSeaVarEXPO [SEA], Weighted AIC: 0.788), with no second or third ranked model for C (Table S5). For clade B the second best model was a birth death model with constant death (DCST) and birth exponentially dependent on the regional South American temperature (BSAMTempVarDCSTEXPO, Weighted AIC: 0.136). African regional temperatures are in the top two ranked models for clades A, D, E. Clade A and D have a pure birth model dependent on African regional temperature (BAFRTempVarEXPO; Weighted AIC: 0.218, A; 0.342, D) as second best preceded by a pure constant birth model (BCST; Weighted AIC: 0.63) for clade A and a time dependent pure birth model (BTimeVarEXPO; Weighted AIC: 0.643) in clade D. The best fitting model for clade E is a pure birth model dependent on African regional temperature (BAFRTempVarEXPO: Weighted AIC: 0.578) followed by a time dependent pure birth (BTimeVarEXPO; Weighted AIC: 0.322) as second-best model (Table S5).

Our GeoHiSSE analyses indicated that *Xylopia* exhibits significant geographic-dependent diversification across different geographic regions and habitat (wet-humid/dry and ultramafic/non-ultramafic). Lineages in Africa, Madagascar, and Central America have higher diversification rates than other regions (geographic-dependent diversification; Tables 3, S6, Fig. 4). Australia + New Guinea however, had a lower diversification rate than other regions. Lineages in the Pacific, Neotropics, and Southeast Asia did not show evidence for geographic-dependent diversification (Tables 3, S6, Fig. 4A).

**Fig. 4.**
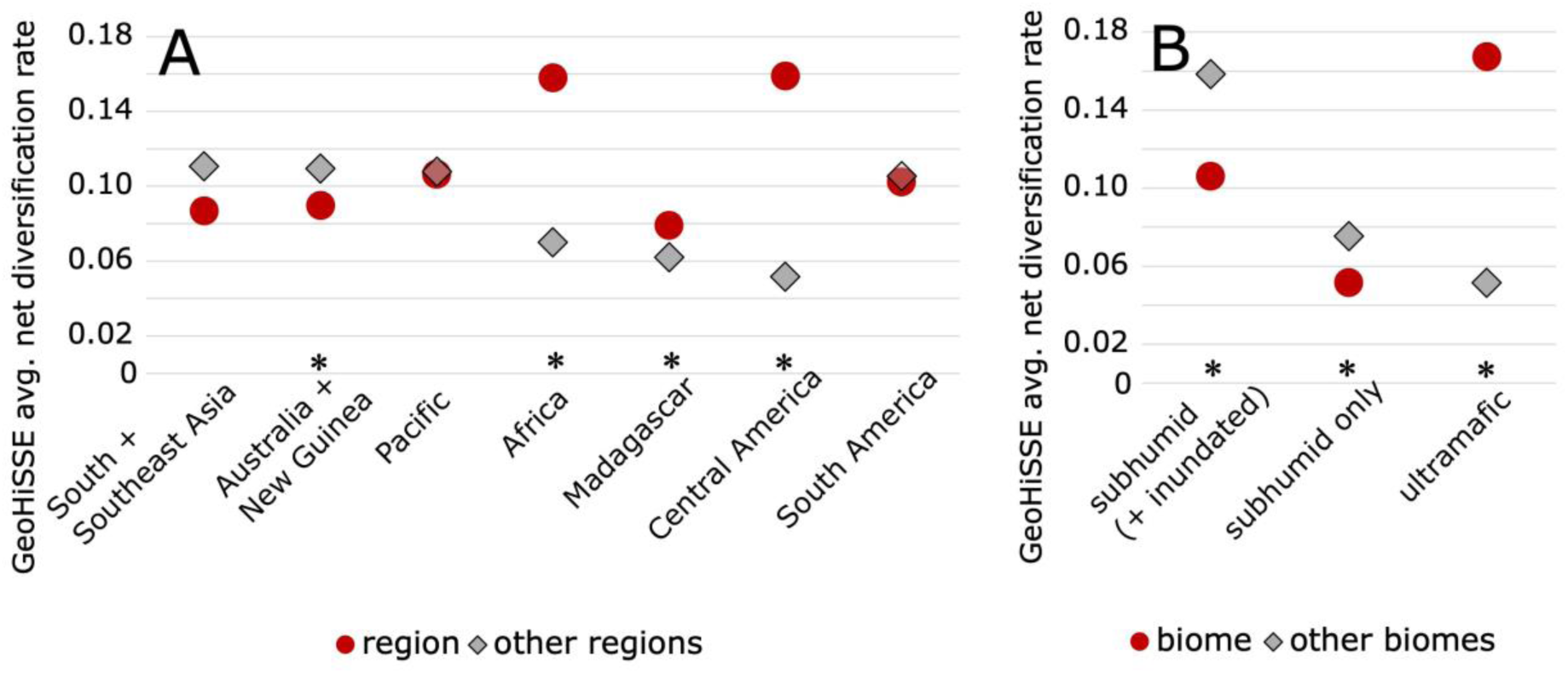
Average net diversification rates for each region and biome based on their respective best-fitting model from GeoHiSSE. Regions and biomes with geographic-dependent diversification as the best-fitting model are indicated by ‘*’.

For humid vs. subhumid lineages, models 22 and 4 were the best-fitting models, both showing geographic-dependent diversification (Tables 3, S7, Fig. 4B). A similar result of geographic-dependent diversification was obtained from our sensitivity test (inundated subhumid taxa coded as ‘humid’) (Table S7). In both cases, subhumid lineages have lower diversification rates than humid rain forest lineages (Table 3). We also show that ultramafic lineages have significantly higher diversification rates than non-ultramafic lineages, based on model 22 as the best-fitting model (Tables 3, S7).

**Table 3.** Best-fitting GeoHiSSE models for each region and biome, based on the highest weighted AIC score. Models with geographic-dependent diversification are highlighted in bold and marked with ‘*’. The average net diversification of each region, biome and their counterpart states are also shown. CID stands for character-independent diversification (i.e., non-geographically dependent).

| Region | Best-fitting model | model specifications | average net diversification rate | average net diversification rate (other state) |
| --- | --- | --- | --- | --- |
| South + Southeast Asia | model 9 | CID—GeoHiSSE extirpation, three hidden rate classes, null model (jump-dispersal forbidden) | 0.0870 | 0.1109 |
| Australia + New Guinea | **model 22\*** | GeoHiSSE, two rate classes, full model (jump-dispersal allowed) | 0.0897 | 0.1095 |
| Pacific | model 9 | CID—GeoHiSSE extirpation, three hidden rate classes, null model (jump-dispersal forbidden) | 0.1065 | 0.1078 |
| Africa | **model 22\*** | GeoHiSSE, two rate classes, full model (jump-dispersal allowed) | 0.1580 | 0.0700 |
| Madagascar | **model 22\*** | GeoHiSSE, two rate classes, full model (jump-dispersal allowed) | 0.0791 | 0.0622 |
| Central America | **model 10\*** | GeoHiSSE extirpation, two hidden rate classes, full model (jump-dispersal forbidden) | 0.1590 | 0.0516 |
| South America | model 3 | CID-GeoHiSSE, three hidden rate classes, null model (jump-dispersal forbidden) | 0.1025 | 0.1055 |
| **Biomes** |  |  |  |  |
| subhumid vs rain forest | **model 22\*** | GeoHiSSE, two rate classes, full model (jump-dispersal allowed) | 0.1063 | 0.1584 |
| subhumid vs rain forest (+ inundated) | **model 22\*** | GeoHiSSE, two rate classes, full model (jump-dispersal allowed) | 0.0517 | 0.0755 |
| ultramafic vs non-ultramafic | **model 22\*** | GeoHiSSE, two rate classes, full model (jump-dispersal allowed) | 0.1674 | 0.0515 |

## Discussion

### Phylogenomic relationships within *Xylopia*

Our phylogenomic results differ somewhat to previous studies of *Xylopia*. Namely, section *Neoxylopia* is no longer sister to section *Rugosperma* as inferred previously based on plastid DNA (Thomas *et al*. 2015; Stull *et al*. 2017). Instead, section *Neoxylopia* forms a grade with other sections. Other discrepancies include the Neotropical *X. peruviana* which belongs to the Neotropical clade of this study, but was shown to be sister instead to African *X. aethiopica* and other Malagasy taxa in Stull et al. (Stull *et al*. 2017). The ST-Verdcourtia clade was sister to the rest of section *Stenoxylopia* in Stull *et al*. (2017) and Thomas *et al*. (2015) but was strongly nested within the section in our study. These differences are likely due to a combination of sampling and different molecular signatures from plastid (of previous studies) and nuclear topologies indicating the presence of reticulate evolution (incomplete lineage sorting or ancient introgression; Guo *et al*. 2018; Stull *et al*. 2023). Incongruences between our concatenated and coalescent topologies are limited to mainly African and Malagasy lineages (and several from Brazil and Sri Lanka, *X. xylantha* and *X. championii* respectively) which may indicate geographic region-specific drivers promoting reticulate evolution. The restriction of these incongruences across shallow nodes of *Xylopia* differs from other Annonaceae (e.g. the Miliuseae tribe) where significant topological conflicts were detected throughout the backbone (Nge *et al*. 2024). Our *Xylopia* crown estimates of *c.* 31.8 Ma in the Oligocene were comparable to that of previous studies despite using different molecular data and calibration points (Thomas *et al*. 2015; Xue *et al*. 2020).

### Diversification rate heterogeneity

Diversification rate heterogeneity detected within *Xylopia* contrasts with gradual accumulation of species and constant diversification of other rain forest lineages. In addition, we provide greater insights into the diversification dynamics of clades within this pantropical genus which were undersampled in previous studies. We show that there is a significant diversification rate increase at crown of the core *Xylopia* clade (sect. *Xylopia* + sect. *Stenoxylopia*). This contrasts with the results of Xue *et al*. (2020) who showed a rate increase at the crown of *Xylopia*, albeit with lower sampling. Higher diversification rates along the core *Xylopia* backbone (from both BAMM and ClaDS) suggests an early burst model of diversification, with near synchronous radiations across continents. Our BAMM results show a single rate shift of increased diversification for *Xylopia*, coinciding with the Oligocene-Miocene boundary (OMB; c. 23 Ma). This period marked an interval of colder temperatures globally, characterized by a major, 400kyr-long glaciation in Antarctica (Mi-1) associated with a 30-40m sea-level drop, likely linked to the combined effects of decreasing atmospheric CO_2_ concentration and favourable orbital parameters (Westerhold *et al*. 2020; Jenny *et al*. 2024). Such forcings likely had environmental consequences at the global scale since biotic turnover and extinction have been noted in several regions across the globe, and interacted with proximal factors such as topographic uplifts (Kamikuri et al. 2005; Deng et al. 2021). In addition, both the extinction of incumbent lineages and increased climatic seasonality opened up niches for the radiation of other lineages during the mid-Miocene (e.g. Nürk et al. 2015; Thode et al. 2019; Nagy 2020; Bacci et al. 2022; Nge et al. 2023). For *Xylopia*, the significant rate shift in diversification during the OMB resulted in the two largest sections in the genus. These two sections (sect. *Xylopia* & sect. *Stenoxylopia*) comprise 55 and 100 species each respectively, representing 92% (155/168) of the total sampled diversity in this study. Biogeographic reconstructions in a companion paper to this one (Johnson et al. unpublished) indicate that the crown radiations of these two sections occurred in Africa, and most likely in rain forest biomes initially. Paleorecords from Africa spanning the OMB recorded a significant decline in tropical forest diversity during this transition (Jacobs *et al*. 2010; Currano *et al*. 2021). The initial diversification of *Xylopia* sect. *Xylopia* and sect. *Stenoxylopia* contrasts to the general floristic response and may represent an example of ecological release and associated radiation, possibly linked to the fragmentation of rain forest habitats during that time spurring allopatric speciation. However, we also acknowledge that there are limited fossil records across Africa from the late Oligocene and early-mid Miocene (Jacobs *et al*. 2010; Couvreur *et al*. 2021). Additional studies are required to reconstruct a more accurate picture of vegetation change during those periods.

The rate-through-time (RTT) curve for *Xylopia* from BAMM showed a sharp increase in speciation at the OMB (c. 23 Ma) consistent with the origin of many plant clades in Africa (Couvreur *et al*. 2021), followed by a decline towards the present. Interestingly, increases in diversification from the OMB was also detected across Annonaceae in general (Fig. 4 in Li et al. 2024). However, the timing of radiations for individual Annonaceae genera and clades are idiosyncratic. Some clades such as Bocageeae have much older radiations than *Xylopia* (Lopes et al. 2024) whereas many other African clades are much younger with recent Pliocene– Pleistocene radiations (Brée *et al*. 2020; Dagallier *et al*. 2023a). The crown radiations of *Uvaria* and *Artabotrys* occurred shortly after the Eocene–Oligocene boundary (c. 33.9 Ma) similar to *Xylopia* (Zhou et al. 2012; Chen et al. 2019), suggesting synchronous radiations of African Annonaceae when other tropical rain forest lineages declined (Couvreur *et al*. 2021). The relatively recent diversification of several Annonaceae genera is not restricted to Africa, but also present in other tropical regions (Erkens et al. 2007; Su and Saunders 2009; Pirie et al. 2018; Xue et al. 2020) and indeed also for other tropical rain forest families (e.g. Richardson et al. 2001; Janssens et al. 2009; Thomas et al. 2012; Koenen et al. 2015; Roalson and Roberts 2016). Speciation declines towards the present from our RTT analyses (BAMM, ClaDS, TESS) may indicate that *Xylopia* and other rain forest lineages have been negatively affected by global cooling and increased seasonality since the Pliocene (Morley 2000; Westerhold *et al*. 2020; Couvreur *et al*. 2021; Jaramillo 2023). Altogether, these findings suggest that the assembly of tropical rain forest diversity is complex and occurred in pulses often in concert with large-scale climatic change, rather than from a constant accumulation of diversity through time.

### Geographic-dependent diversification

We showed that different environmental drivers have affected the diversification of *Xylopia* clades within each region, based on our RPANDA analyses. Not surprisingly, *Xylopia* lineages in the tectonically dynamic and complex Asia-Pacific area (with many islands) had speciation and diversification rates correlated most with past sea-level fluctuations. These sea-level changes may have promoted lineage divergence from repeated cycles of fragmentation and re-connection of landmasses, essentially acting as species-pumps (Yang *et al*. 2013; Li and Li 2018). The different diversification modes (constant rate and time-dependent) for two Malagasy clades further demonstrate the complexity and heterogeneity of evolutionary responses across lineages, even within the same region. Only the African clade (ST-acutiflora + ST-odoratissima) showed regional temperature-dependent diversification, in contrast to the finding of Li *et al*. (2024) that temperature-dependent diversification correlates with speciation and extinction rates across all of Annonaceae. However, these differences may be scale-dependent with their study focusing on broader family-level diversification dynamics. Nevertheless, we and others (Tiatragul *et al*. 2023) showed that there is greater nuance in regional dynamics than realised in studies that have only used global-scale environmental data for these models. In contrast to our results, studies on another clade of African Annonaceae (Monodoreae tribe; Dagallier *et al*. 2023a) showed that orogeny across the African continent (particularly in eastern Africa from early Miocene onwards) was the best explanatory factor for changes in speciation rates in that particular tribe, thus reinforcing the notion of group-specific responses to different environmental drivers even within the same region or plant family.

Our study showed significant differences in diversification rates among regions (from GeoHiSSE analyses) for *Xylopia* despite synchronous radiations across them. Regions with younger clades (Africa, Madagascar, and Central America) had significantly higher diversification rates compared with older clades in other regions. This finding is thus in agreement with others showing for many organismal groups that diversification rates are often scaled to time, with fastest rates found in younger clades (Henao Diaz *et al*. 2019; Wiens 2024). The largest African radiation (ST-odoratissima + ST-acutiflora) is younger (*c.* 10 Ma) than similar sized counterparts in Asia (ST-malayana, *c.* 12.5 Ma) and South America (*c.* 12 Ma), albeit the confidence intervals of these estimates overlap. Higher diversification rates in Africa for *Xylopia* go against the common narrative of lower diversification rates (due to greater extinction) and lower species richness of African clades compared to other tropical regions (Plana 2004; Couvreur 2015; Hagen *et al*. 2021). Higher rates in Africa for *Xylopia* might be linked to biotic traits such as smaller leaf size as an adaptation to more xeric environments, responding to past climatic changes across the continent (Johnson et al. unpublished).

Interestingly, all five Malagasy *Xylopia* clades radiated or diverged from 5-7 Ma (late Miocene), with significant lag time from their stem divergence suggesting a synchronous response to a common driver in the region. The onset of heavy seasonal rain regionally during the late Miocene is linked to the establishment of the Indian monsoons during that time (Molnar *et al*. 1993; Buerki *et al*. 2013), driven by a number of paleogeographic factors particularly orogeny in eastern Africa and the complete emergence of the Arabian plate (Couvreur *et al*. 2021; Tardif *et al*. 2023). The expansion of rain forest coinciding with onset of the Indian monsoonal regime has resulted in radiations of many Malagasy biotic assemblages during the late Miocene (8–10 Ma), evident from our study and many others (Zhou *et al*. 2012; Armstrong *et al*. 2014; Strijk *et al*. 2014; Gamisch and Comes 2019; Salmona *et al*. 2019; Godfrey *et al*. 2020; Farminhão *et al*. 2021; Skema *et al*. 2023). These radiations were not mentioned in previous reviews on the evolutionary assembly of Malagasy groups as they had focused on origins (stem divergence) and colonisation events instead of radiations (Buerki *et al*. 2013; Antonelli *et al*. 2022). Shifting patterns of habitat connectivity and fragmentation from escarpment erosion across eastern Madagascar may also play a role in these *Xylopia* radiations as has been shown for other Malagasy plants (Liu *et al*. 2024).

Australia + New Guinea was the only region with significantly lower diversification rates for *Xylopia* compared to other regions globally. In Australia, we attribute this to the limited extent of rain forest on the continent and the contraction of mesic habitats following Miocene aridification (Byrne *et al*. 2011). Indeed, many endemic Australian plant and animal clades show speciation declines and extinction towards the present in response to severe aridification (e.g. Renner *et al*. 2020; Toussaint *et al*. 2022; Nge *et al*. 2023; Nge *et al*. 2025). That diversification rates in *Xylopia* are significantly lower in New Guinea is surprising given it has the world’s richest island flora (Cámara-Leret *et al*. 2020). However, the island is not particularly species-rich in terms of *Xylopia* species (Johnson and Murray 2023). Indeed Madagascar, an island about 25% smaller than New Guinea, contains nearly four times as many species, 30 versus 8 respectively (Johnson and Murray 2020; Johnson and Murray 2023), despite recent taxonomic revisions on both regions. In addition, it may be premature to determine the diversification dynamics of rain forest lineages in general in New Guinea due to lack of taxonomic studies targeting the region (Su and Saunders 2009; Cámara-Leret *et al*. 2020; Johnson and Murray 2023; Ezedin 2024). Most available studies on the diversification of the New Guinean biota have focused on uplift-driven radiations (i.e., non-rainforest lineages; Toussaint *et al*. 2014; Oliver *et al*. 2017; Toussaint *et al*. 2021; Roycroft *et al*. 2022).

The relatively low diversification rates of the three early diverging sections of *Xylopia* and coupled with their ‘broom and handle’ divergence signature suggests the clades have undergone substantial extinction, followed by subsequent re-diversification in the Miocene (Crisp and Cook 2009). Two of these clades (sect. *Neoxylopia* and sect. *Ancistropetala*) are African in origin (Johnson et al. unpublished), and likely experienced extinction during the Oligocene–Miocene boundary congruent with the wider flora in response to climate cooling and increased aridification across the continent (Couvreur 2015; Currano *et al*. 2021). However, we did not detect any mass extinction events from our CoMET analyses across *Xylopia* similar to another Annonaceae study on African Piptostigmateae (Brée *et al*. 2020). It is possible that these extinctions are specific to these two sections and hence were not detected in our genus-wide analyses. The diversification signatures of sect. *Rugosperma* are less clear as half of the extant diversity (4 spp.) were not sampled. Further sampling on this section would shed light on its diversification dynamics: the crown radiation of this clade is substantially older than the other Asian clade of the genus (ST-malayana + ST-peekelii clade).

### Diversification in subhumid environments

The negative association between transitions into subhumid habitats and diversification rates for *Xylopia* reflects that these transitions consist largely of lineages with only one or few extant species. Indeed, only less than half (4/19) of these transitions have resulted in radiations (Johnson et al. unpublished). It could be that certain traits have allowed *Xylopia* lineages to transition and persist (lower extinction rates) in drier environments but not speciate (Johnson et al. unpublished). Our findings are in contrast to others, showing no significant difference (Veranso-Libalah *et al*. 2018; Zizka *et al*. 2020) or significantly higher diversification rates in subhumid compared to rain forest environments for other plant groups (Simon *et al*. 2009; Bouchenak-Khelladi *et al*. 2010). These differences highlight the clade-specific nature of diversification following transitions out of rain forest habitats. In *Xylopia*, the two largest subhumid radiations (both in core *Xylopia*) occurred in Africa and might be linked to the continent having the largest area of seasonally dry forests and savanna compared to other continents (Pennington *et al*. 2018) (Johnson et al. unpublished). The relatively high number of transition events into subhumid conditions for *Xylopia* compared to other Annonaceae genera corroborate findings of Boyko and Vasconcelos (2024) that speciation and extinction (turnover) rates are correlated with transition rates across biomes. Indeed, *Xylopia* is one of the few genera/clades detected to have a significant positive diversification rate shift in Annonaceae (Xue *et al*. 2020). Thus, it is not transitions into subhumid habitats *per se* that are linked to diversification but rather the number and rate of transition events. Conversely, higher diversification rates for *Xylopia* in tropical rain forests points to the more dynamic nature of this biome, with higher turnover, *in situ* speciation, and floristic exchange (migration) compared to other tropical biomes (Pennington and Dick 2004; Pennington *et al*. 2018)(Johnson et al. unpublished).

### Diversification on ultramafic substrates

Higher diversification rates in ultramafic sites compared to other habitats suggests that lineages in *Xylopia* were able to successfully overcome and exploit extreme niches in stressful environments. Indeed, there were two independent *Xylopia* radiations onto ultramafic substrates – one in New Caledonia and another smaller radiation in Cuba (Johnson et al. unpublished). The specific growing environment of ultramafic substrates requires plants to physiologically deal with high concentrations of metals, hence limiting many plant groups from establishing in this particular environment (Pillon *et al*. 2010; Garnica-Díaz *et al*. 2023). *Xylopia* has the highest proportion of ultramafic species (*c.* 10%) compared to other Annonaceae genera (Johnson et al. unpublished), suggesting that certain clades in the genus might have been predisposed (exapted) to colonise and radiate in ultramafic environments (Johnson et al. unpublished)(Pillon *et al*. 2019). As in *Xylopia*, transitions to growing on ultramafic substrates have resulted in higher diversification rates for other plant groups (Condamine *et al*. 2017b; González Gutiérrez *et al*. 2023), but this pattern is by no means universal (Pérez-Calle *et al*. 2024). These findings indicate more broadly that specialists are not always doomed to extinction (‘evolutionary dead ends’), and can even be a more successful evolutionary strategy than generalists (Vamosi *et al*. 2014; Day *et al*. 2016). Focusing on trait-dependent diversification, specifically traits that have allowed *Xylopia* species to transition, persist, and radiate in ultramafic and other non-rain forest habitats along with dispersals across continents are promising avenues for future research (Onstein *et al*. 2019).

## Conclusion

In this study, we present a densely-sampled species level phylogenomic temporal framework for the pantropical genus *Xylopia* to investigate diversification dynamics of tropical rain forests at a global scale. We show that despite near synchronous radiations across tropical regions, there is significant diversification rate heterogeneity along the backbone of core *Xylopia*, and across different regions, with clades responding to different environmental conditions. Transitions to novel (non-rain forest) environments do not always lead to elevated diversification rates, with higher rates noted for ultramafic but not subhumid clades. Regional diversification models indicate sea-level changes as the most important environmental driver for Asian, Australian, Pacific, and Neotropical clades (including archipelagos of Central America), i.e., regions with many islands that have experienced repeated cycles of fragmentation and reconnection of different landmasses through time. Whereas diversification of the largest *Xylopia* clade on the African continent and Madagascar (ST-odoratissima + ST-acutiflora) was best explained by regional temperature changes. A companion paper (Johnson et al. unpublished) showed that the timing and number of dispersal events from Africa to other continents also differ for clades in *Xylopia* yet still allowing the genus to achieve a pantropical distribution in the Miocene. Overall, our studies support the idea that rain forests across different regions are dynamic, complex and non-uniform in their evolutionary histories. Further studies on other rain forest lineages can shed light on whether these region-specific diversification and biogeographic patterns in *Xylopia* are also applicable to the wider regional biotas more generally.

## Acknowledgements

This project has received funding from the European Research Council (ERC) under the European Union’s Horizon 2020 research and innovation program (GLOBAL project; grant agreement No. 865787) to TLPC.

We acknowledge the ISO 9001 certified IRD i-Trop HPC (South Green Platform) at IRD Montpellier for providing HPC resources that have contributed to the phylogenetic results reported within this paper.

## Data availability

The paired fastq sequences included in this study are available in Genbank SRA under Bioproject number PRJNA1265870. Scripts used for analyses are available on Github:

