## Supplementary materials for "Synchronous Miocene radiations and geographic-dependent diversification of pantropical *Xylopia* (Annonaceae)"

**Fig. S1.** Phylogenetic tree (RAxML) with outgroups (uploaded separately as pdf).

**Fig. S2.** BEAST dated divergence tree (uploaded separately as pdf).

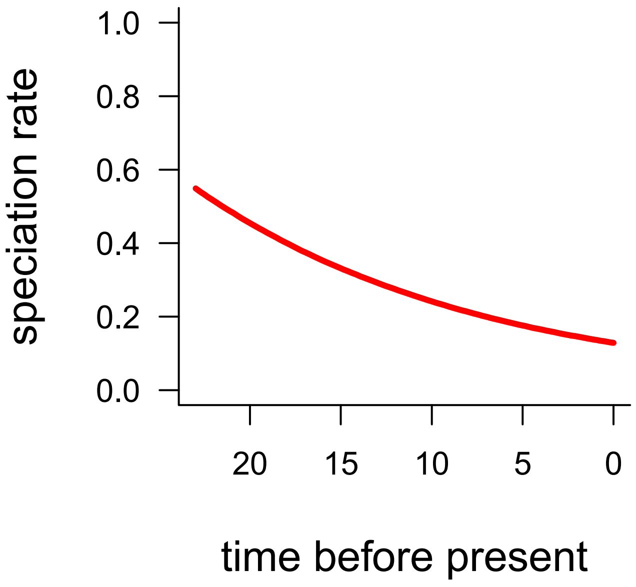


**Fig. S3.** Rates-through-time plot for the core *Xylopia* clade (Sections *Xylopia* + *Stenoxylopia*; node 172) from BAMM.

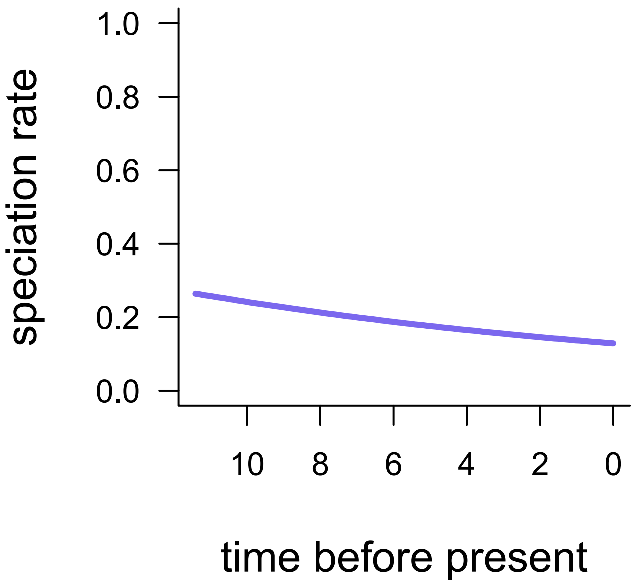


**Fig. S4.** Rates-through-time plot for the Neotropical *Xylopia* subclade (node 218) from BAMM.

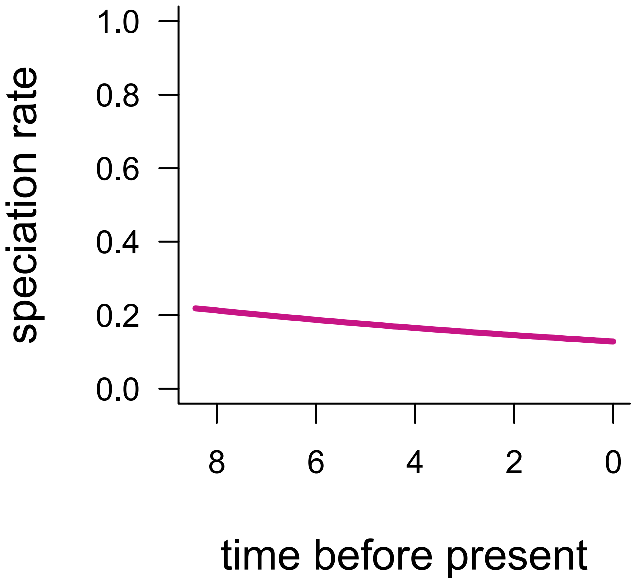


**Fig. S5.** Rates-through-time plot for a Madagascar clade of *Xylopia* subclade (node 273) from BAMM.

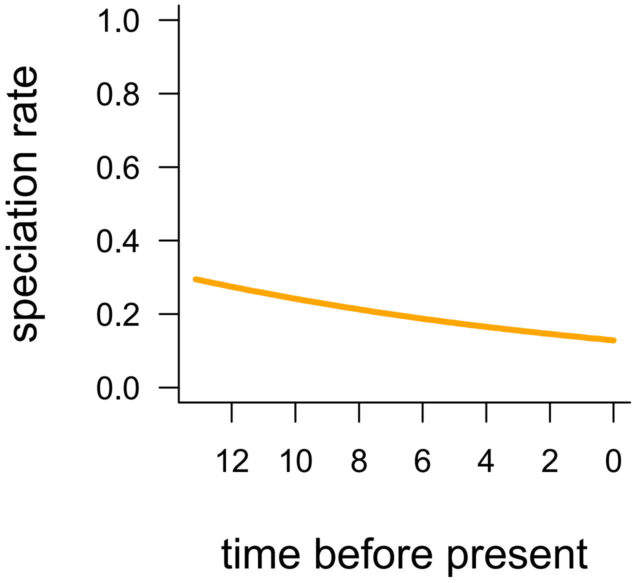


**Fig. S6.** Rates-through-time plot for the Asian + Australia + New Guinea + Pacific clade of *Xylopia* subclade (node 230) from BAMM.

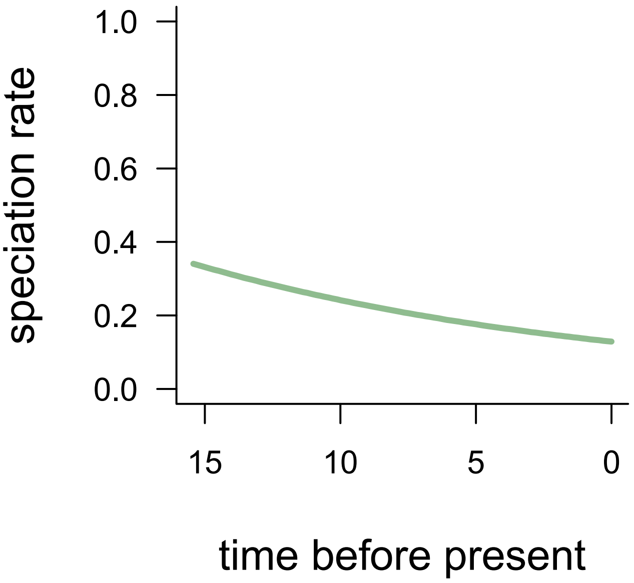


**Fig. S7.** Rates-through-time plot for the main African clade of *Xylopia* (node 174) from BAMM.

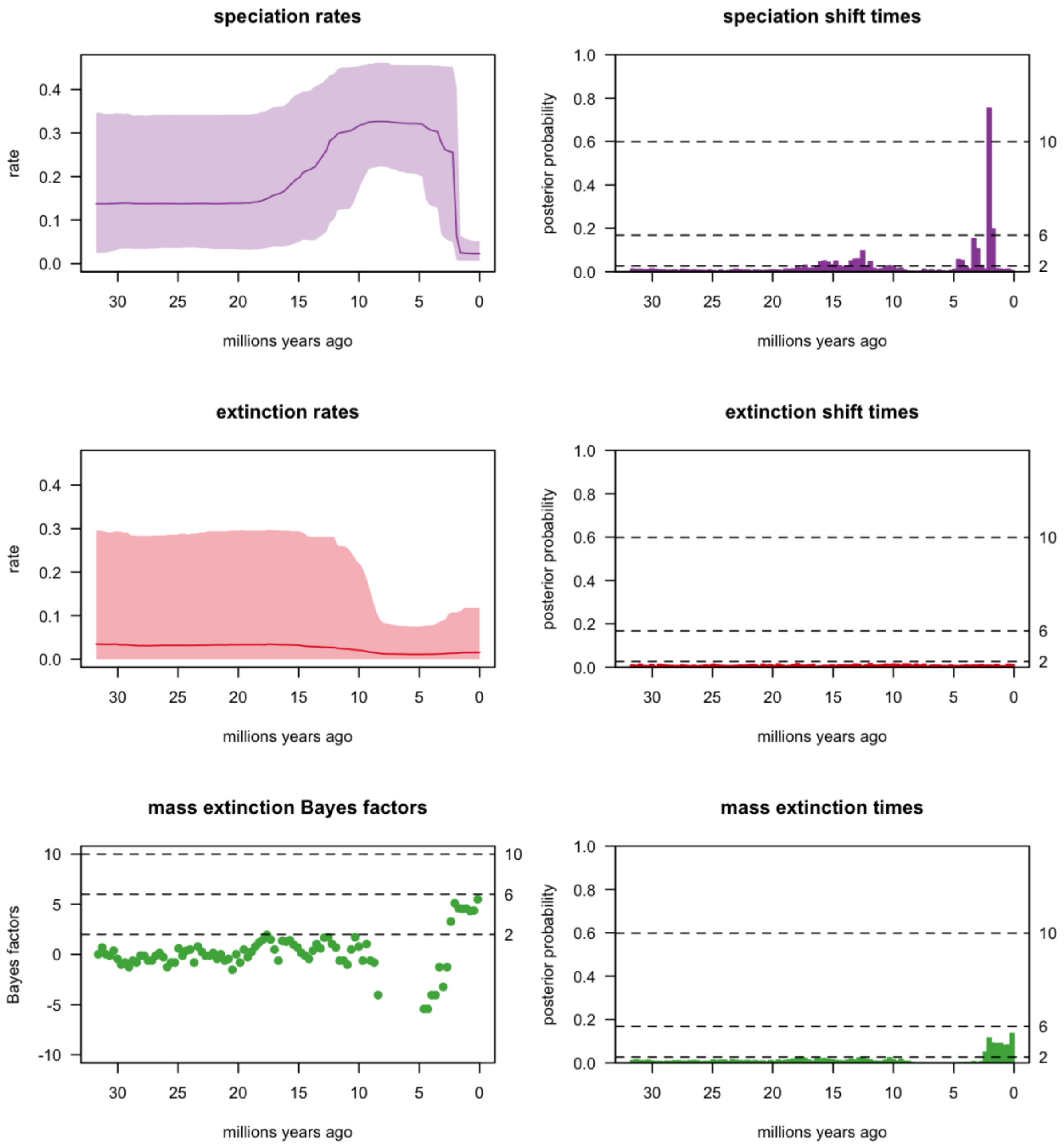


**Fig. S8.** Summary figures from TESS and CoMET showing rates-through-time (speciation and extinction), posterior probability of speciation/extinction shifts, and mass extinctions. These analyses were completed based on the specified prior of 50% survival probability for mass extinction events.

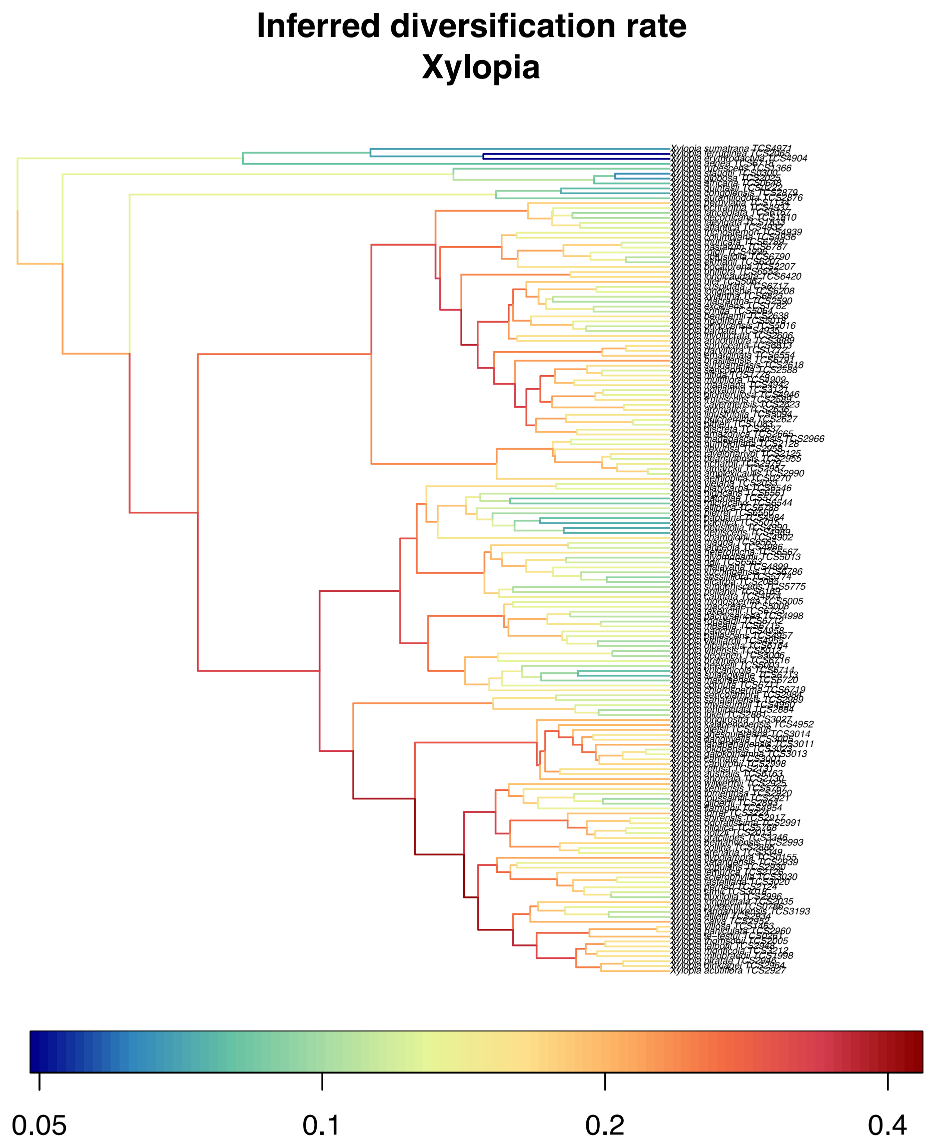


**Fig. S9.** Inferred diversification rate mapped onto the *Xylopia* phylogeny, based on the ClaDS analysis.

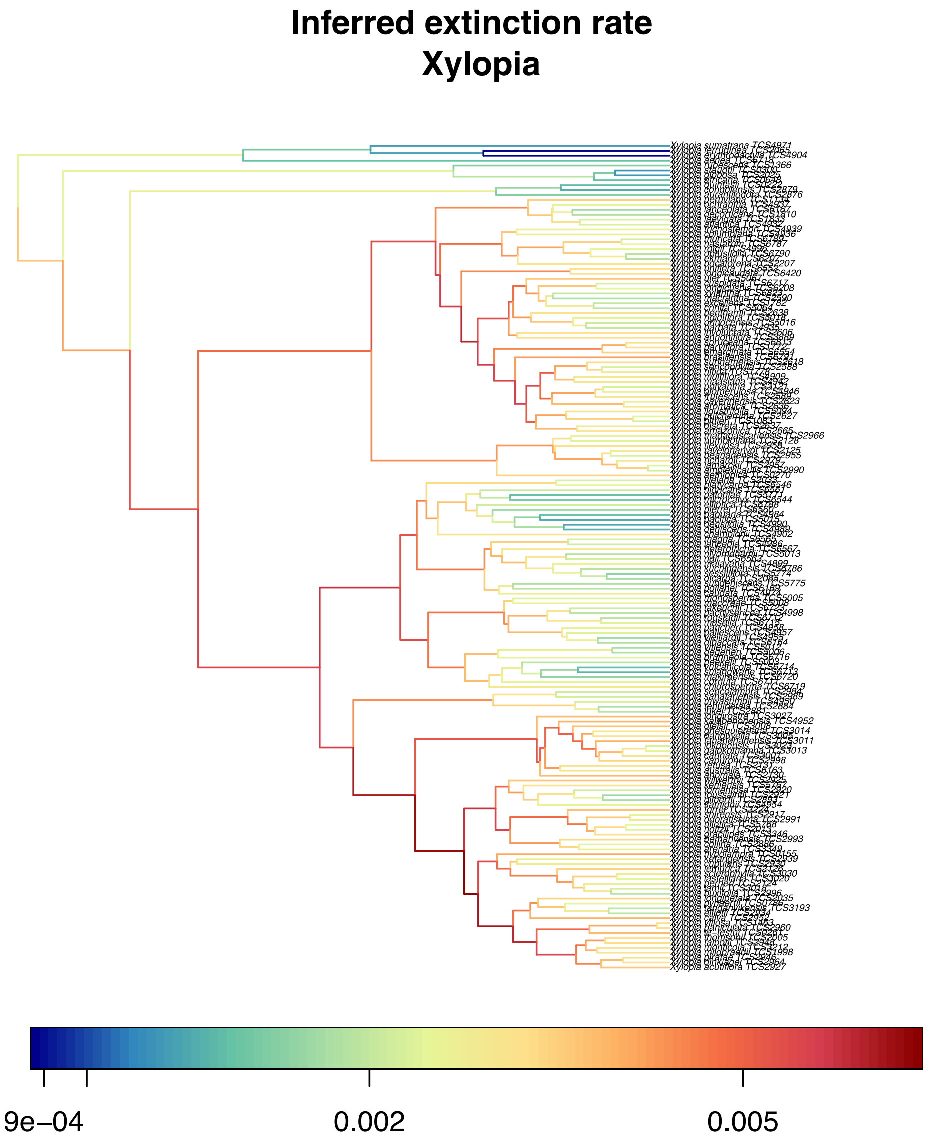


**Fig. S10.** Inferred extinction rates mapped onto the *Xylopia* phylogeny, based on the ClaDS analysis.

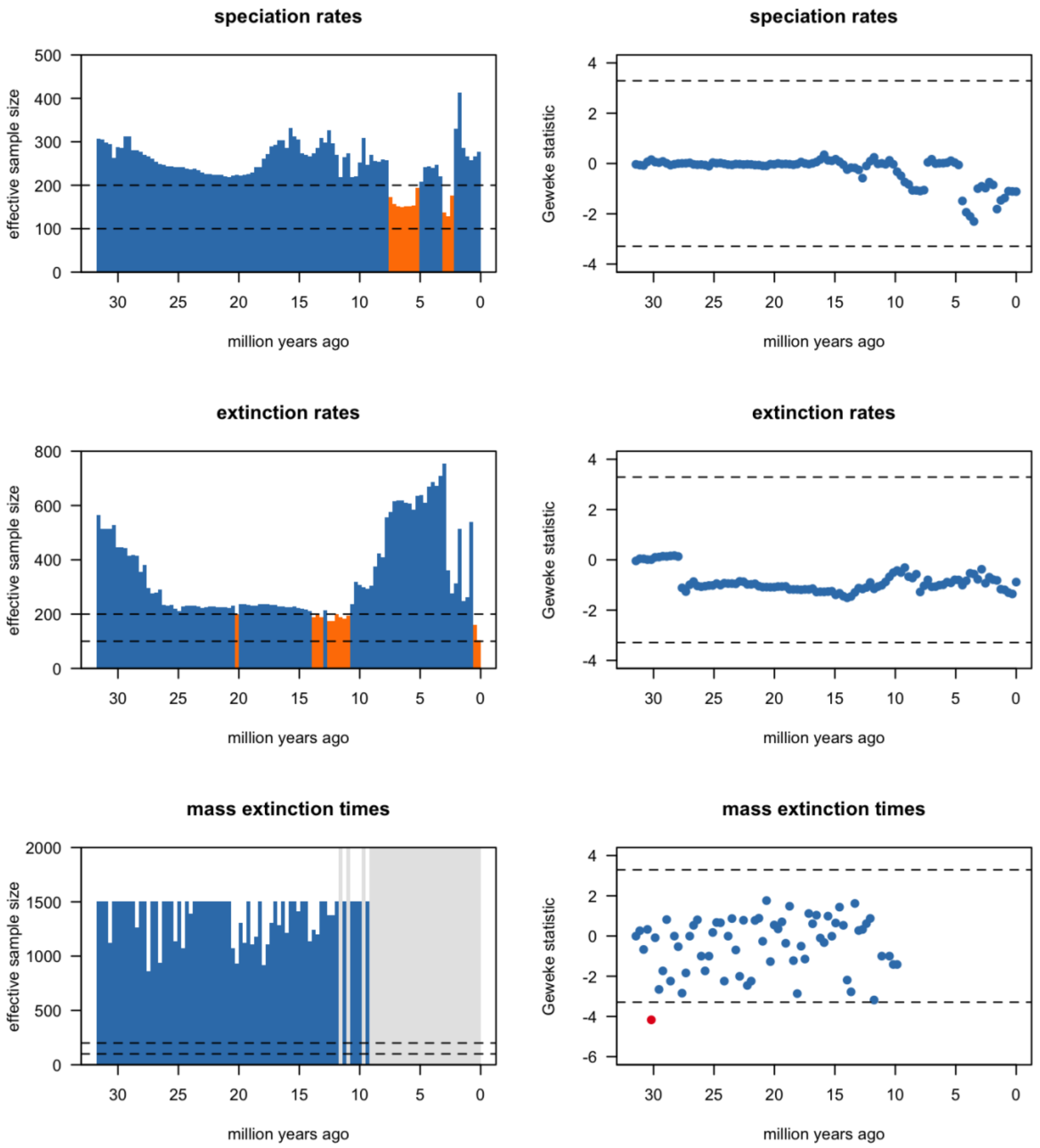


**Fig. S11.** Effective sample size (ESS) for TESS and CoMET analyses with convergence achieved at ESS > 200. These analyses were completed based on the specified prior of 20% survival probability for mass extinction events.

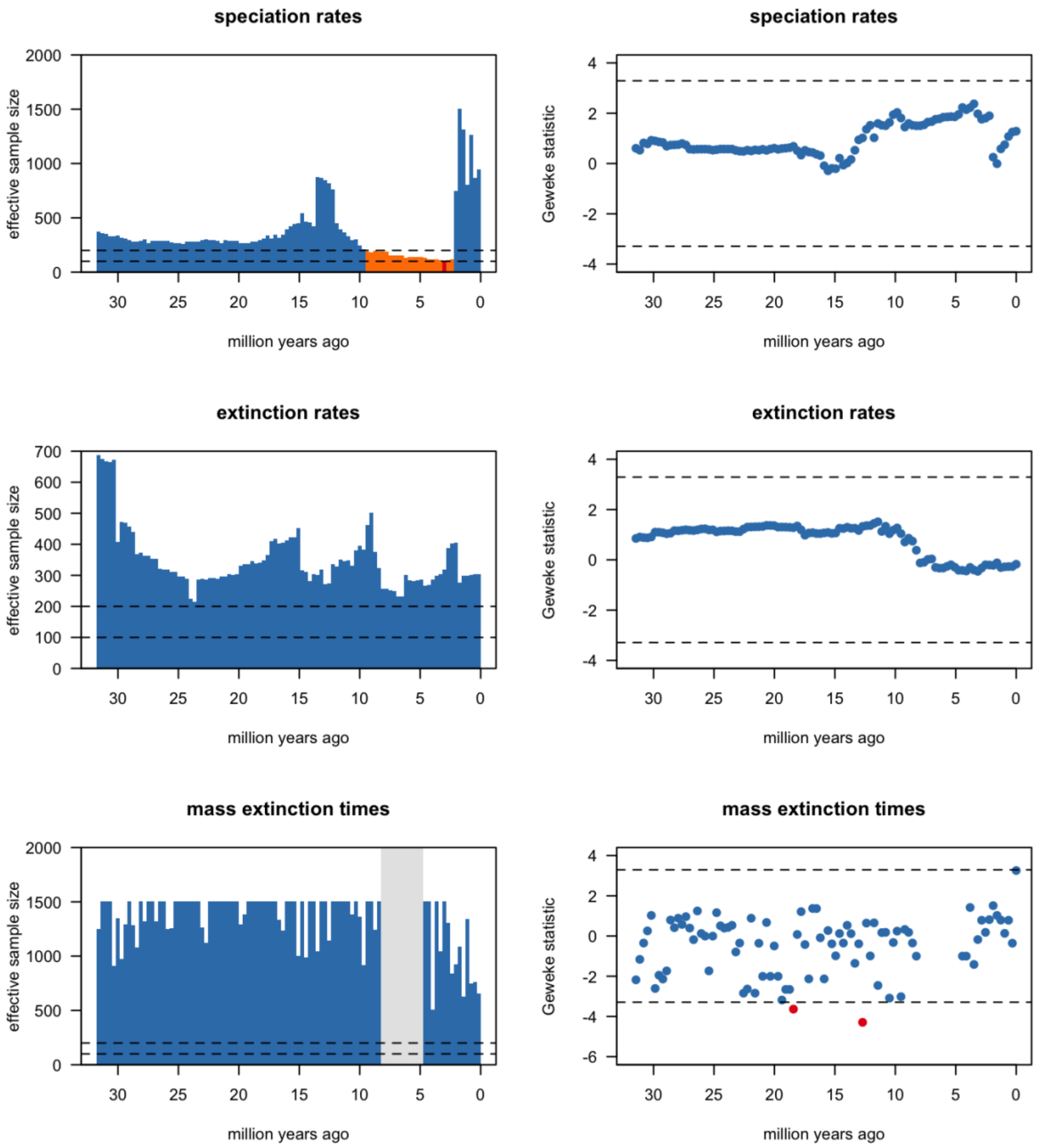


**Fig. S12.** Effective sample size (ESS) for TESS and CoMET analyses with convergence achieved at ESS > 200. These analyses were completed based on the specified prior of 50% survival probability for mass extinction events.

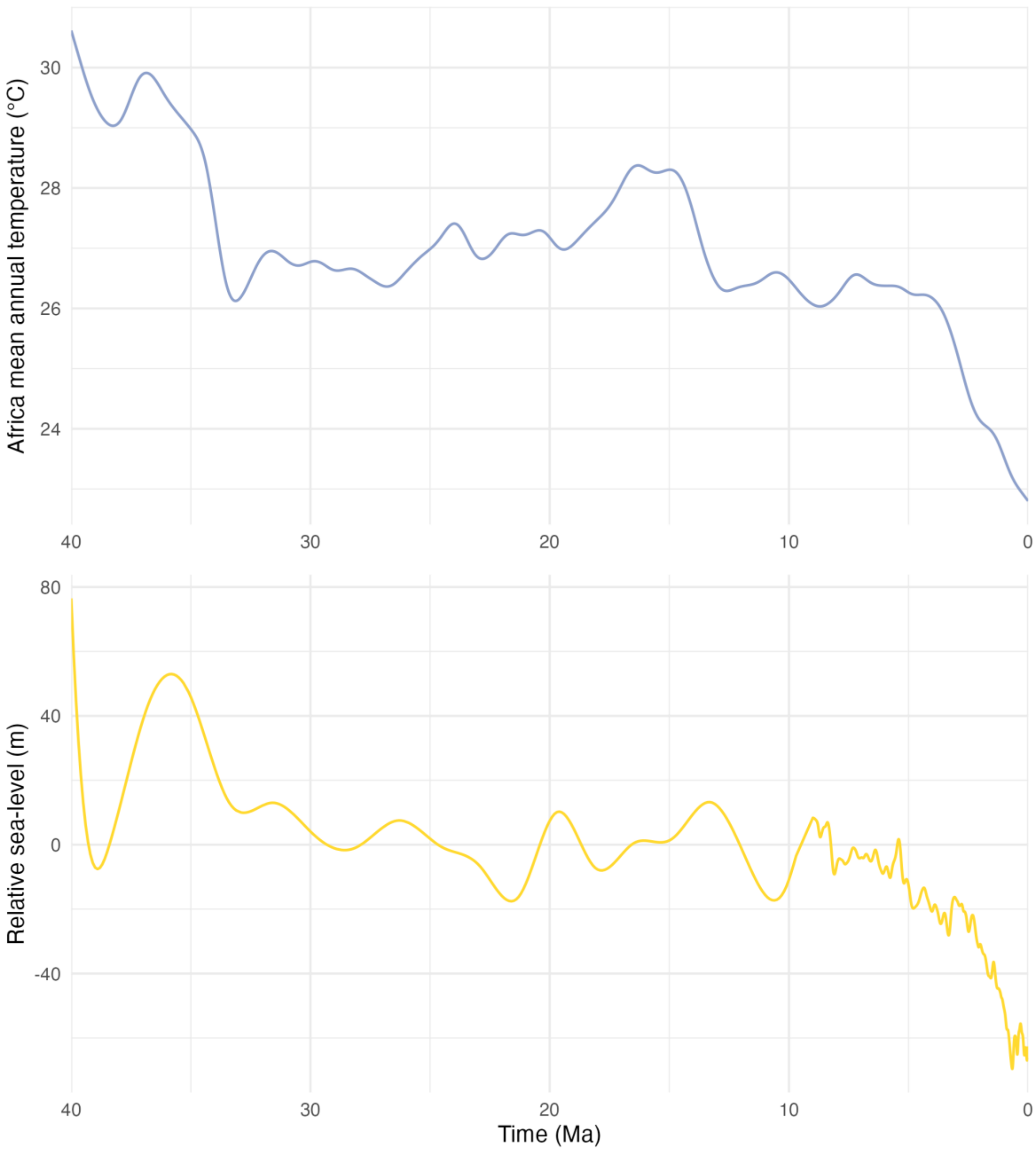


**Fig. S13.** Temperature and sea level curves used in the RPANDA models, with region-specific African mean annual temperature and relative sea-level (global).
